# Base editing strategy allows insertion of the A673T mutation in APP gene to prevent the development of Alzheimer’s disease

**DOI:** 10.1101/2020.10.27.357830

**Authors:** Antoine Guyon, Joël Rousseau, Francis-Gabriel Bégin, Tom Bertin, Gabriel Lamothe, Jacques P. Tremblay

## Abstract

Amyloid precursor protein (APP), a membrane protein mostly found in neurons, is preferentially cut by the α-secretase enzyme, however, abnormal cleavage by β-secretase leads to the formation of β-amyloid peptide plaque in the brains of Alzheimer’s patients. Genome analysis of an Icelandic population that did not appear to show symptoms of Alzheimer’s at advanced age led to the discovery of the A673T mutation, reducing β-secretase cleavage by 40%. We hypothesized that the insertion of this mutation in a patient’s genome could be an effective and sustainable method to slow down or prevent the progression of familial and sporadic forms of Alzheimer’s disease. We have thus modified the APP gene in HEK293T cells and in SH-SY5Y neuroblastoma using a Cas9n-deaminase enzyme, which changes a cytosine into a thymine, thus converting the alanine codon to a threonine. Several Cas9n-deaminase variants were tested to compare their efficiency of conversion. The results were characterized and quantified by deep sequencing. We successfully modified the APP gene in up to 56.7% of the HEK293T cells. Our approach aimed to attest to the efficiency of base editing in the development of treatments against genetic diseases as well as provide a new strategy for the treatment of Alzheimer’s.

## INTRODUCTION

Alzheimer’s disease (AD) is one of the most well-known neurological illnesses due to its prevalence in the elderly population and the lack of effective treatment. It is responsible for 70 % of the reported cases of dementia, currently totaling about 47.5 million cases worldwide with an estimated progression to 75.6 million in 2030 according to the World Health Organization. Patients are afflicted with memory loss, temporal and spatial confusion, and have difficulty planning tasks^(1)^. The onset and development of this disease can be attributed to the accumulation of amyloid plaques between neurons. This eventually results in the death of the patients^(2)^. Unfortunately, a treatment capable of attenuating the accumulation of amyloid plaques has yet to be created. These plaques are the result of the undesirable cleavage profile of the amyloid β precursor protein (APP).

APP is a membrane protein that is preferentially cleaved by α-secretase. However, cutting of the protein by the β-secretase (BACE1) leads to the formation of amyloid-β peptides. These peptides form aggregates which accumulate as plaques between neurons in the brains of patients afflicted with Alzheimer’s ^(3, 4)^.

Numerous mutations in the APP gene have been shown to favor β-secretase cutting and thus the accumulation of plaques leading to Familial Alzheimer’s Disease (FAD). However, it was discovered that an APP gene variant (Icelandic mutation A673T) present in a Scandinavian population reduces β-secretase cleavage in the APP by 40% and also reduces the β-peptide aggregation^(5)(6)^. This mutation of the alanine codon into a threonine codon is due to the modification of a single base pair in exon 16 which. Individuals carrying this mutation present almost no accumulation of amyloid-β peptides in their brain even at 95 years of age. Moreover, the A673T mutation has been linked to a greater life expectancy as carriers of this mutation are 50% more likely to reach the age of 85 when compared to non-carriers. Further substantiating this claim, Kero et al.’s identified a 104 year old person carrying the 60 A637T gene who later died with little β-amyloid pathology^(7),(8)^.

This evidence suggests that a therapy based on transmitting this mutation to AD patients or genetically susceptible individuals would be beneficial to prolong their quality of life by slowing down the development of the disease. To this end, we are proposing the use of the Clustered Regularly Interspaced Short Palindromic Repeats (CRISPR) system to introduce this A673T mutation. While this system was originally used by bacteria such as *Streptococcus pyogenes* (Sp)^(9)^ and *Staphylococcus aureus* (Sa)^(10)^ as an immune system to resist bacteriophages, researchers worldwide have successfully co-opted it for their own purposes.

The original CRISPR system used a CRISPR-associated protein (Cas) and a sgRNA to selectively cleave a specific nucleic acid sequence. However, Dr Liu et al. have recently published their development of Cas9 fusion proteins (i.e. BE3 & Target-AID) that they have called base editors ^(11)^. Their fusion proteins contain a cytidine deaminase enzyme and a Cas9 nickase (Cas9n) linked by an amino acid sequence. The final result is a fusion protein that allows for a direct, programmable, and irreversible conversion of a C:G base pair into T:A pair without inducing a Double Strand Break (DSB). The Cas9n fusion protein forms a complex with a single guide RNA (sgRNA) and relies on the later’s sequence to target and attach to a specified DNA complementary sequence. The formation of this protein/RNA/DNA complex liberates an R-Loop, which exposes a small section of single stranded DNA (approximately five nucleotides). All the cytosines present in this area are deaminated and thus converted to uracil, resulting in an intermediate G:U. The cell then engages its base excision repair mechanism to solve the G:U mismatch. This is initiated by the excision of uracil by uracil N-glycosylase (UNG). To protect this intermediate, Liu’s group fused a uracil glycosylase inhibitor (UGI) to the Cas9n C-terminal, increasing the conversion of the G:U to A:U and finally A:T by 50%. The combination of these elements forms the complex BE3 (Base Editing Version 3) (Figure 1). This technique was subsequently improved by the same group ^(12–14)^ with the release of the BE4 system.

**Figure 1:**
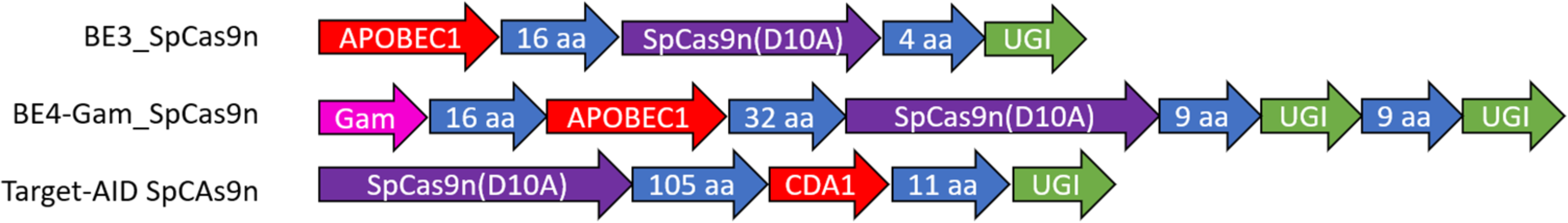
Structures of BE3, BE4 and Target-AID variants. Various cytidine deaminases (APOBEC1, YE1_APOBEC1 or Target-AID (CDA1) were fused with various Cas9 nickases (SpCas9n_VQR_, SaCas9n_KKH_, SpCas9n_EQR_).

One limitation of this system is the dependency the Cas9 protein has on its associated protospacer adjacent motif (PAM). For the Cas9 to induce a DSB in its target sequence, said sequence must contain a pre-defined set of nucleotides, to which the Cas9 will first bind^(15)^. Only after the Cas9 interacts with this sequence can it induce a DNA DSB at a specific site recognized by the sgRNA. As a result, researchers are often limited in their selection of an adequate Cas9 based on the presence of a specific PAM. Since the base editing system relies on a Cas9 for targeting, it is likewise constrained by the presence of an adequate PAM.

In the present study, we used the CRISPR/Cas9 base editing method to mutate the APP gene, allowing a seamless and efficient A673T editing in human cells. We tested several Cas9/deaminase variants in order to find the most efficient enzyme to induce the Icelandic mutation.

## RESULTS

### Deaminase design

BE3_SpCas9n (henceforth designated as BE3) was the original SpCas9n deaminase made by Komor et al. (11) (Figure 1). It contains the APOBEC1 deaminase, a 16 amino acids linker, the SpCas9 nickase (i.e., a SpCas9n containing a D10A mutation to prevent the cut of one DNA strand), a second 4 amino acids liker and the uracil DNA glycosylase inhibitor (UGI). This version of the enzyme recognized a NGG or NGA PAM sequence. Unfortunately, this constraint meant that it could not be used in our experiment since there was no NGG PAM near the codon 673 in exon 16 of the APP gene. As such, it was necessary to use a new base editor to induce the Cytosine to Thymine conversion required for the creation of the A673T mutation.

As shown in Figure 2, the antisense DNA strand was targeted using three different BE3. These base editors differed in their Cas9n enzymes with each possessing different PAMs: NGAN (for SpCas9n_VQR_) ^(16)^, NGAG (for SpCas9n_EQR_) ^(16)^, or AGAGAT (for SaCas9n_KKH_) ^(17)^. This was done to induce the deamination of the cytidine in position 2 (C2) in the editing window in the antisense strand into a thymine (T), thereby changing the alanine codon (GCA) into a threonine codon (ACA) in the sense strand i.e., the A673T mutation.

**Figure 2:**
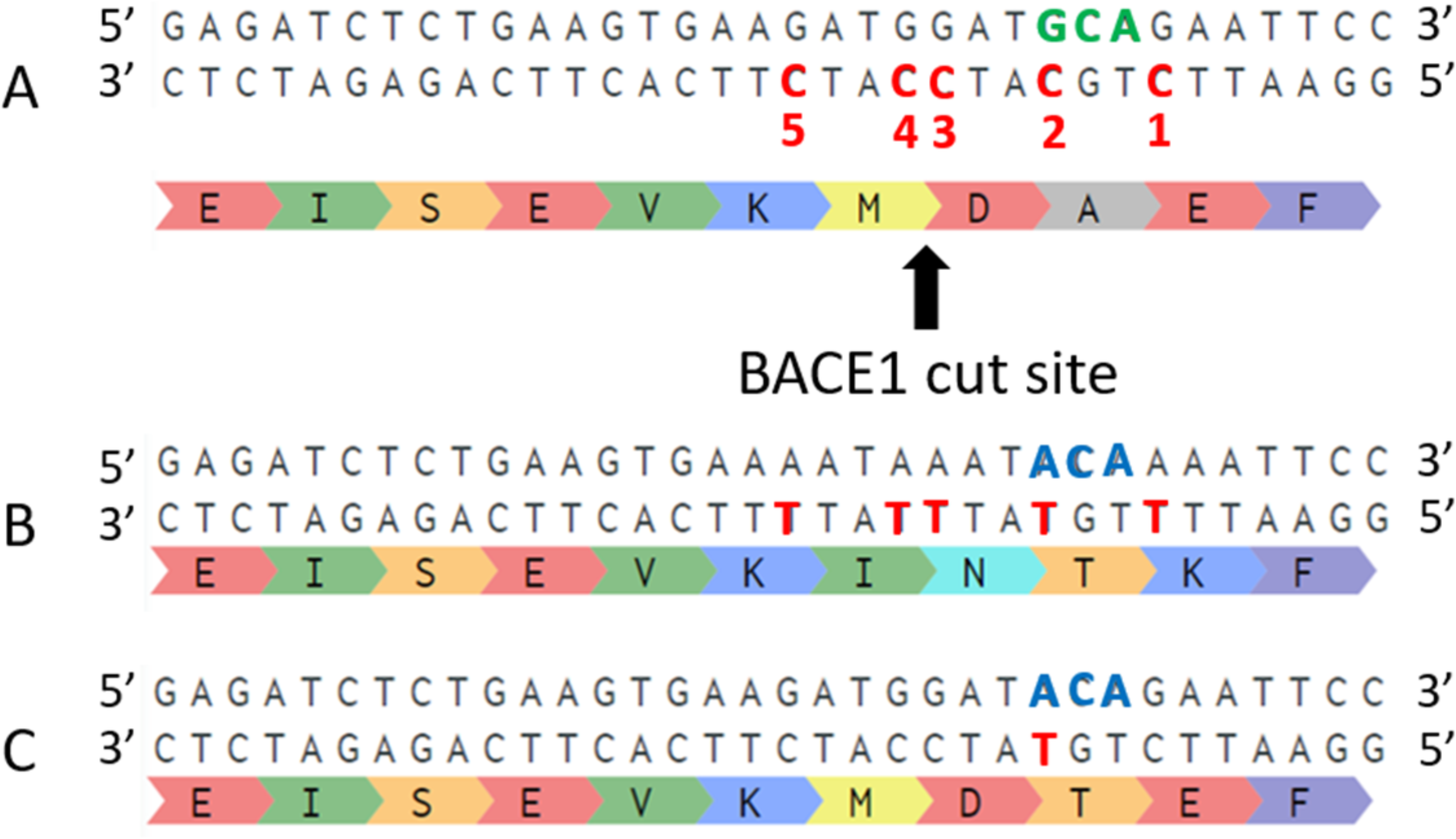
Amino acid modifications produced by base editing of the various cytidines. In A, part of exon 16 of the wild type APP gene is illustrated. In the antisense strand, there are 5 cytidines in red (C1 to C5), which are potentially in the editing window of each base editor. The 673-alanine codon is in green. In B, the sequence of the APP gene in which all five cytidines in the targeting window are deaminated into thymines resulting in the change of four amino acids since the cytidine to thymine modification of the C5 nucleotide is a silent mutation that does not change the resulting amino acid. In C, the sequence of the APP gene when only cytidine C2 has been deaminated changing the alanine codon into a threonine codon (in blue).

Mutant SpCas9 nucleases have previously been produced wit to react with alternative PAMs ^(16)^. Of them, Cas9n_VQR_ and Cas9n_EQR_ were selected for the purposes of this study, The VQR variant has been shown to robustly bind to sequences with NGAN PAMs while the EQR variant is more specific and thus anneals to an NGAG PAMs. These mutant Cas9 molecules were used as a base to create the following base editors. BE3_SpCas9n_VQR_, BE3_SpCas9n_EQR_, BE3_SaCas9n_KKH_, BE4-Gam_SpCas9n, YE1-BE3_SpCas9n and Target-AID SpCas9n were purchased on Addgene. BE4-Gam_SpCas9n_VQR_, BE4-Gam_SpCas9n_EQR_, YE1-BE3_SpCas9n_VQR_, YE1-BE3_SpCas9n_EQR_, YE1-BE3_SaCas9n_KKH_, Target-AID SpCas9n_VQR_, Target-AID SpCas9n_EQR_, Target-AID SaCas9n_KKH_ were made in our laboratory. Several sgRNA lengths, from 17 bp to 22 bp, were tested in order to influence the conversion window (list available in supplemental results Table S1). One or two sgRNA copies were inserted in a modified pBSU6 plasmid.

### APP deamination: A673T editing

Deep sequencing was used to determine the percentage of deamination (i.e., a cytosine changed into a thymine) obtained for each cytosine (C1 to C5) present in the target window of the deaminase (analysis examples available in supplemental data Table S2).

The editing efficiency in HEK293T cells was greater than in SH-SY5Y cells throughout the study (Figure 3A). The three variants of Cas9 used with the APOBEC1 deaminase (BE3_SpCas9n_EQR_, BE3_SpCas9n_VQR_, BE3_SaCas9n_KKH_) showed a similar editing profile, however, SH-SY5Y cells only demonstrated ~40% of the editing found in HEK293T cells (Figure 3B for HEK293T and Figure S1 for SH-SY5Y). BE3_SpCas9n_VQR_ exhibited the highest deamination rate. Inclusion of two copies of sgRNA in the plasmid slightly increased the targeted deamination (Figure 3C). As illustrated in Figure 3D, the best deamination percentages of the C2 nucleotide using APOBEC1 were obtained with the BE3_SpCas9n_VQR_ enzyme with a sgRNA targeting 20 nucleotides in both cell models (SH-SY5Y available in supplemental results Figure S2). C1 and C3 nucleotides were deaminated more frequently than the C2 nucleotide with every base editor using the APOBEC1 enzyme (present in all the BE3 and BE4 constructs). From these experiments, it was concluded that the SpCas9n_EQR_ and the SaCas9n_KKH_ base editor variants were showcasing poor deamination rates and were greatly less effective than the BE3_SpCas9n_VQR_ variant (results available in supplementary results Figure S3). Thus, the BE3_SpCas9n_EQR_ and SaCas9n_KKH_ were not used in the subsequent experiments.

**Figure 3:**
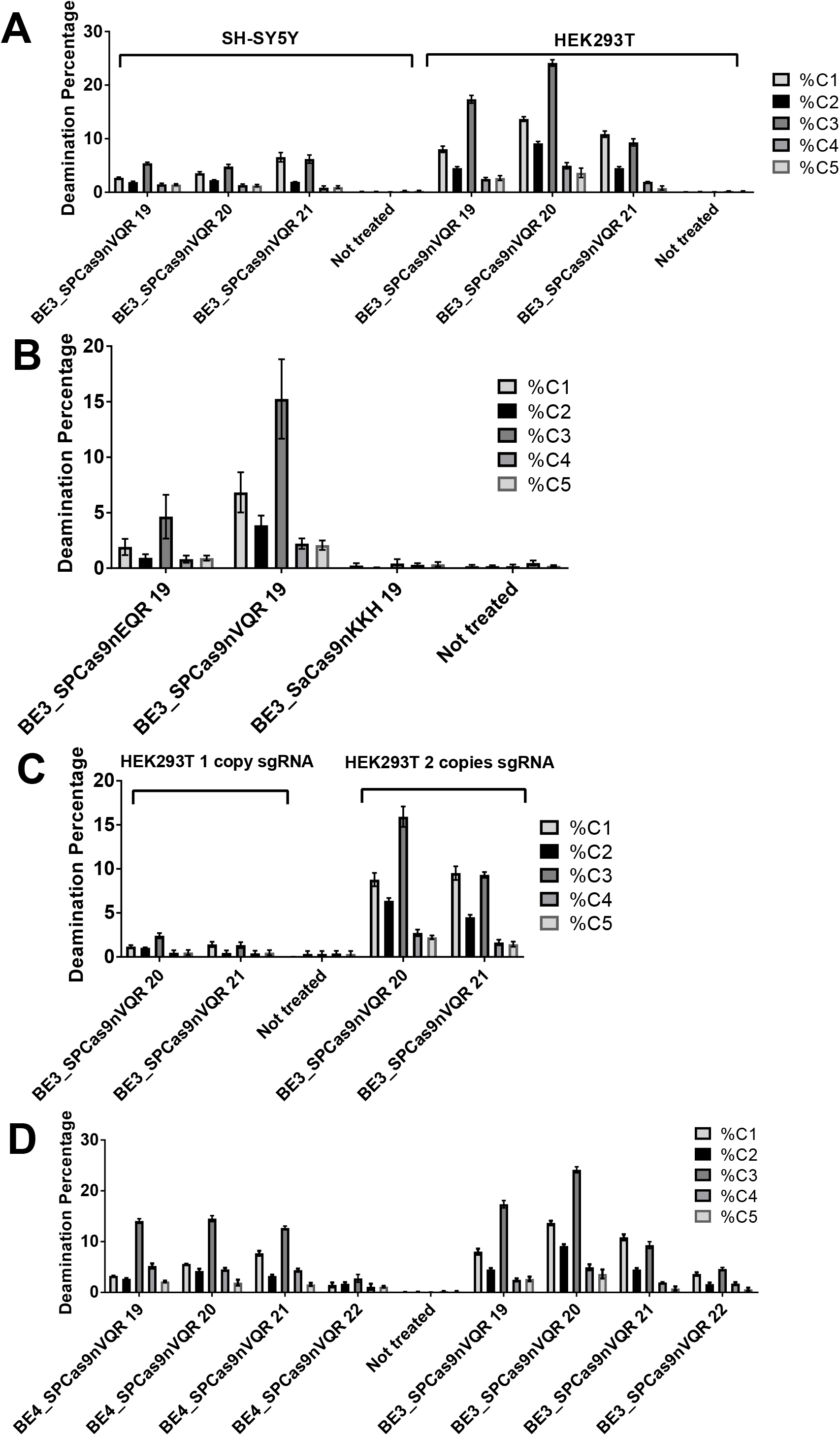
Percentages of cytidine deamination produced by various enzymes and sgRNAs. Plasmids coding for the various base editors and for one or two sgRNA were transfected in HEK293T and SH-SY5Y cells. The number of nucleotides targeted by the sgRNAs is indicated after the name of the enzymes. DNA was extracted 3 days post transfection, APP exon 16 was PCR amplified and sent for deep sequencing. In **A**, difference between SH-SY5Y and HEK2093T editing efficacy is shown with BE3_SpCas9n_VQR_. In **B**, BE3_SpCas9n_EQR_, BE3_SpCas9n_VQR_, BE3_SaCas9n_KKH_ enzymes were tested in HEK293T cells. The figure illustrates the means +/− SEM (n=4). In **C**, the comparison between the use of one copy sgRNA versus two copies during base editor transfection in HEK293T cells. In **D**, the BE4_SpCas9n_VQR_ and BE3_SpCas9n_VQR_ enzymes were tested in HEK293T cells. SH-SY5Y results available in complementary results S1 and S2.

When the first experiments were performed, the conversion window was quite wide. That is, the targeted cytidine (C2) was not deaminated frequently enough compared to the other cytidines. In response, new cytidine deaminase enzymes were designed (YE1-BE3_SpCas9n_VQR_, YE1-BE3_SpCas9n_EQR_, YE1-BE3_SaCas9n_KKH_, Target-AIDSpCas9n_VQR_, Target-AIDSpCas9n_EQR_, Target-AIDSaCas9n_KKH_). The enzymes were tested with sgRNAs targeting 17, 18, 19, or 20 nucleotides this time. Among these new cytidine deaminase enzymes, the Target-AID-SpCas9n_VQR_ deaminated a higher percentage of cytidines, especially C1 and C2, resulting in a narrower target window (Figure 4A for SH-SY5Y / Figure S4 for HEK293T cells). Using a sgRNA targeting either 17 or 18 bp, C2 was preferentially deaminated with statistically significant differences. But the highest deamination of the C2 nucleotide was obtained with a sgRNA binding with 19 nucleotides. A similar deamination profile was observed in the SH-SY5Y and HEK293T cells. A higher deamination rate was noted in SH-SY5Y for the first time in the study, indicating that Target-AID SpCas9n_VQR_ could work better in these cells.

**Figure 4:**
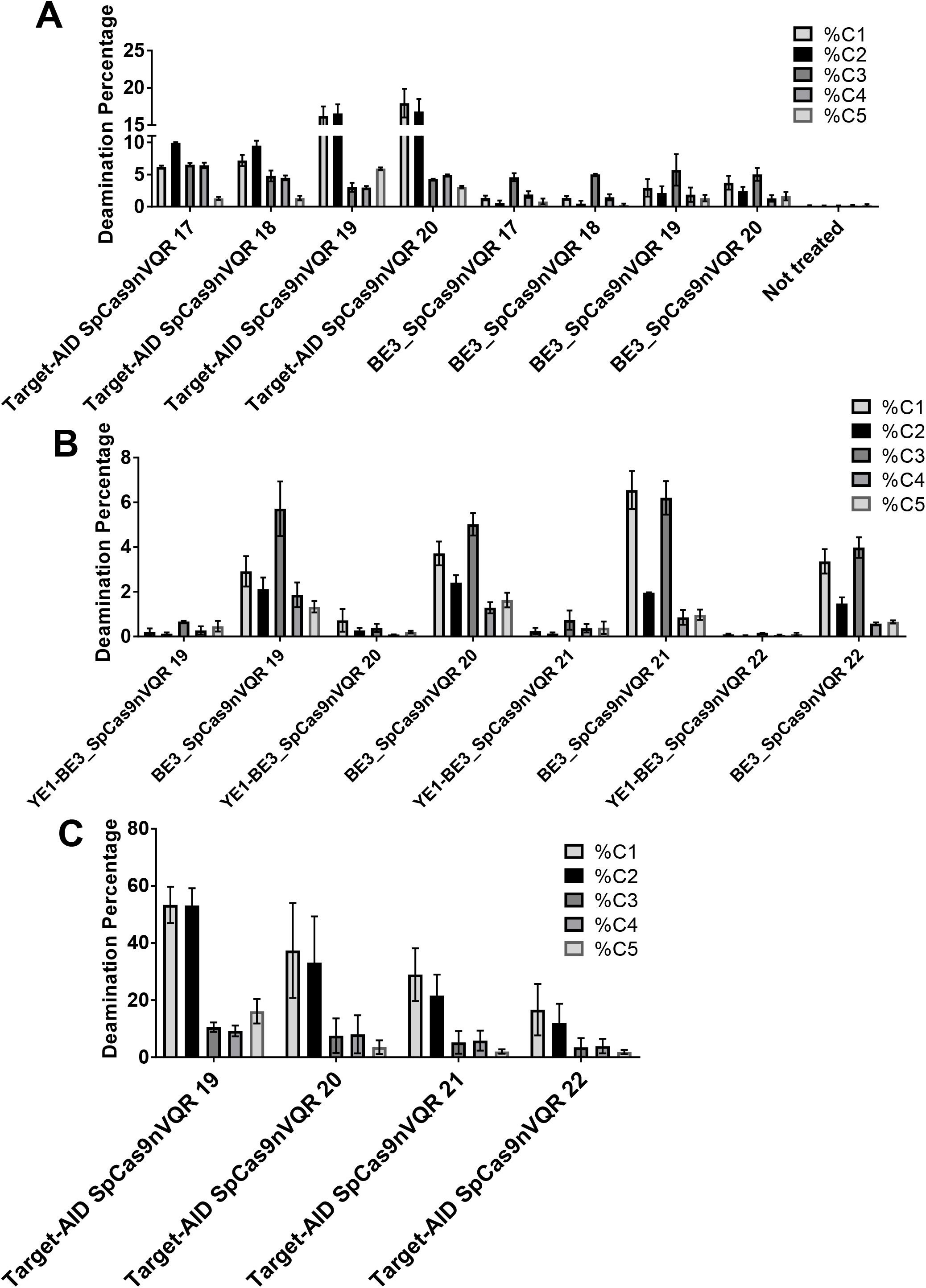
Deamination efficiencies using various Cas9n-deaminases and sgRNAs targeting various numbers of nucleotides. In **A**, the difference of deamination in SH-SY5Y of cytidines C1 to C5 produced by the Target-AID-SpCas9n_VQR_ and BE3_SpCas9n_VQR_ enzymes and two copies of a sgRNA targeting 17 to 20 nucleotides. Error bars are mean with SEM (n=3). In **B**, difference between YE1-BE3_SpCas9n_VQR_ and BE3_SpCas9n_VQR_ in SH-SY5Y cells. The figure illustrates the means +/− SEM (n=4). In **C**, the percentage of deamination in HEK293T cells transfected with Target-AID-SpCas9n_VQR_ and two copies of a sgRNA targeting 17 to 20 nucleotides after optimization. The figure illustrates the means +/− SEM (n=5).

The YE1-BE3 variant was also tested. This variant contains two mutations W90Y and R126E in the APOBEC1 gene and has previously been reported to create a narrower conversion window^(12)^. The enzyme was tested with sgRNAs targeting 19, 20, 21, or 22 nucleotides. This variant converted an insignificant percentage of cytidine C2 into thymine in exon 16 of the APP gene with the three Cas9 variants (expected for SpCas9n_EQR_ and SaCas9KKH) (Figure 4B for SH-SY5Y / Figure S5 for HEK293T).

The experiment was repeated with Target-AID-SpCas9n_VQR_ and with the BE3_SpCas9n_VQR_ combined with sgRNA targeting 19, 20, 21, or 22 nucleotides. The highest percentages of deamination of the nucleotides C1 and C2 were obtained with the Target-AID-SpCas9n_VQR_ used with sgRNAs targeting 19 nucleotides (Figure 4C). We therefore decided to select this variant for the further steps.

### Aβ peptides concentration decreases in APP SH-SY5Y cell lines

The next experiment was based on the work previously performed in our laboratory, which found that the A673T mutation was responsible for reducing the Aβ peptide concentration, which would otherwise had been produced by an APP gene containing a FAD mutation^(8)^. The best FAD mutation responding to the treatment being the London mutation V717I (expected decrease in Table S3). The aim was to deaminate plasmids coding for APP containing a FAD mutation and attesting an Aβ40/42 concentration decrease as proof of principle. Different SH-SY5Y cell lines that constitutively expressed wild-type APP, V717I APP or C1+C2 mutations of APP (E674K, A673T) were produced through lentiviral transduction. These cells were subsequently transfected with a lentivirus plasmid EF1-Target-AID-SpCas9n_VQR_-2U6 gRNA19. Interestingly, as observed in Figure 5A, deaminated SH-SY5Y APP cell lines demonstrated different deamination profiles. Indeed, C1 was not as frequently deaminated as C2. The two cell lines transfected respectively had 6.64% C2 for SH-WT and 7.57% for SH-V717I. A reduction in the Aβ40 and Aβ42 peptides secreted in the culture medium was observed (Figure 5B). Culture medium containing cells overexpressing WT APP experienced a reduction of 19.9% Aβ40 and 6.7% for Aβ42. London mutation V717I APP cell line demonstrated a 26.4% Aβ40 and 31.8% Aβ42 reduction, and the C1+C2 cell line showcased a 43.8% Aβ40 and 52.7% Aβ42 reduction in peptides compared to WT APP.

**Figure 5:**
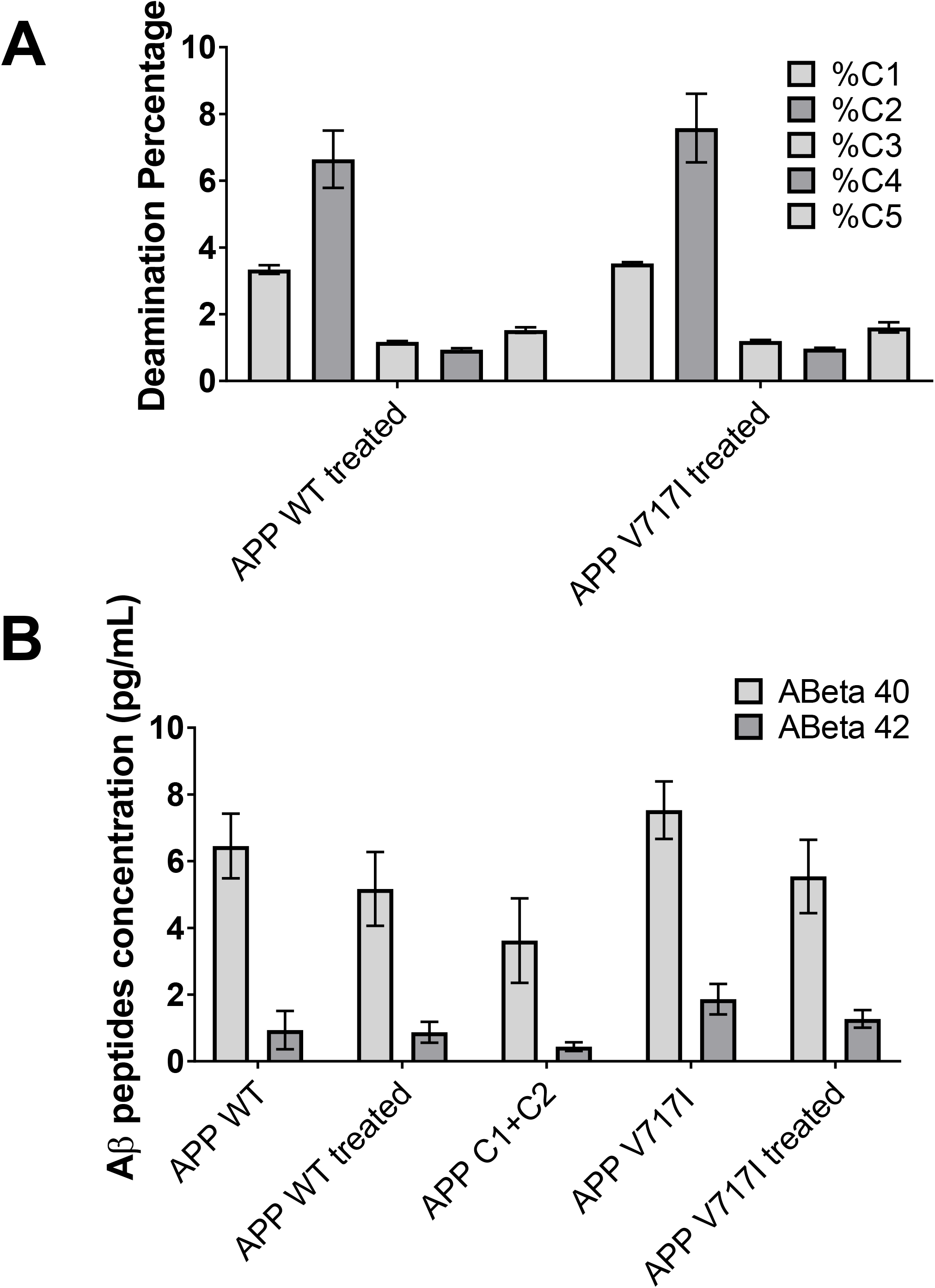
Concentration of Aβ42 peptides in the cell supernatant of SH-SY5Y APP cells. SH-SY5Y cells were transfected with a lentivirus plasmid containing Target-AID-SpCas9n_VQR_ and two copies of the sgRNA targeting 19 nucleotides. In **A**, Target-AID-SpCas9n_VQR_ deamination profile in SH-SY5Y cell lines for Wild-type and V717I APP. In **B**, Aβ42 peptides concentration in SH-SY5Y supernatant. The figure illustrates the means +/− SEM (n=3).

## DISCUSSION

In this study, we have focused on the development of a promising base editing technique that was designed to introduce the Icelandic A673T mutation in patients susceptible to AD as a means of granting them protection towards the accumulation of beta peptide plaques. The beneficial effects of this mutation were demonstrated through its introduction into SH-SY5Y cells lines containing either the wild-type APP gene, the C1+C2 mutated APP or a variant of this gene containing the London mutation V717I^(8)^. Treated cell lines WT and London showcased respectively 19-26% reduction in Aβ40 peptides and 6.7-31.8% in Aβ42 peptides respectively, which is promising despite a low transfection efficacy for our future *in vivo* studies. C1+C2 cell line was created in order to attest if the deamination profile obtained with our ideal base editor was creating a better or worse reduction of Aβ peptides. Surprisingly the reduction obtained was better than a common A673T mutation with 43.8% Aβ40 and 52.7% Aβ42 (Table S3). The process to determine which mutation endowed the greatest protection against FAD required numerous screenings using various purchased and lab-made base editors.

The CRISPR/Cas9-deaminase complex that we determined to be the most efficient preferentially deaminated the targeted cytosine but also weakly deaminated the other four nearby cytosines. The ideal choice for a base editor is often locus dependent; looking at our study, SpCas9_VQR_ was the most effective of the three Cas9 variants. This could be different in another gene. We also obtained an explanation for the reduced deamination of the C2 nucleotide with APOBEC1. Indeed Dr. Komor’s team informed us this cytidine deaminase was inhibited when a G was preceding the target C (our C2 target is preceded by a G). BE3 and BE4 base editors were indeed deaminating C1 and C3 predominantly, which was not beneficial towards our proposed treatment. Target-AID, on the other hand, showed a much higher deamination rate with a narrower conversion window around C1 and C2. Target-AID-SpCas9n_VQR_ which presented the highest deamination rate was a new base editor designed in our lab using Target-AID-SpCas9n as a model. Recent base editors have been developed since the beginning of this study such as XBe3 and evoCDA1-BE4max^(18, 19)^. Unfortunately, the first one was using an APOBEC1 deaminase and the other seemed to have a wider conversion window than our Target-Aid using a CDA1 construct. Since there are several C in our area of interest, this obliged us to select the narrowest editing window possible. The reduced production of the Aβ42 and Aβ40 peptides observed in the cell supernatant suggests that the additional cytosine mutations did not alter the protective effect of the A673T mutation. It would be beneficial to investigate the effects these unintended mutations might engender *in vivo*. It will be necessary to verify whether these additional mutations affect the aggregation of the peptides or have any other detrimental effects. We hope that the development of the PRIME editing technology is going to help us to limit the modifications only to the C2 nucleotide ^(20)^.

The off targets from base editing are extremely small with no Cs modified outside of the conversion window in our case. While indels may be more common in gene therapies that involve Cas9-mediated cleavage or the recently designed prime editing with the PE3 technique using a second sgRNA to induce a second nick, base editing approaches are primarily subject to off targeting in the conversion window. An *in-silico* investigation using Benchling.com interface for our gRNA off targets demonstrated no notable off target events (Figure S6). When looking at one mismatch situations, no off targets were found. A possible off-target site was predicted only when our search was widened to two mismatches, however this off-target was located in a non-coding DNA sequence.

For this type of gene editing, one of the most important factors is the safe and efficient delivery of the therapeutic agents. For the *in vivo* experiments and for an eventual clinical application, a dual AAV vector delivery system may prove the only option to introduce the cytidine deaminase transgene in the neurons. In fact, a single virus could not package a base editor given its size. Base editing could be used to treat many hereditary diseases, even though some specific optimizations will be required to adjust the target in the genome.

## MATERIAL AND METHODS

### Deaminase variants description and construction

The enzymes that we have tested are the following:

1. **BE3_SpCas9n**_**VQR**_, produced by Kleinstiver et al.(16), is a variant of BE3, which contains an SpCas9n protein with 3 mutated amino acids D1135V/R1335Q/T1337R. The gene for this enzyme was available at Addgene Inc. as pBK-VQR-BE3 (#85171).
2. **BE3_SpCas9n**_**EQR**_, also produced by Kleinstiver et al. (16), is another variant of BE3, which also contains an SpCas9n protein with 3 mutated amino acids D1135E/R1335Q/T1337R. The gene for this enzyme was available at Addgene Inc. as pBK-EQR-BE3 (#85172).
3. **BE3_SaCas9n**_**KKH**_ is a variant of BE3, also produced by Kleinstiver et al (17), which contains an **Sa**Cas9n protein (from Staphylococcus aureus) with 3 mutant codons E782K/N968K/R1015H. The gene for this enzyme was available at Addgene Inc. as pJL-SaKKH-BE3 (#85170).
4. **YE1-BE3_SpCas9n** is a construct variant of BE3, which contains 2 mutant codons (W90Y/R126E) in the rAPOBEC1 sequence. The gene for this enzyme was available at Addgene Inc. as pBK-YE1-BE3 (#85174). We used this plasmid to construct all other variants containing the YE1 rAPOBEC1 deaminase.
5. **YE1-BE3_SpCas9n**_**VQR**_ is a variant of the YE1-BE3_SpCas9n made in our laboratory. The rAPOBEC1 deaminase sequence in plasmid pBK-VQR-BE3 was replaced with restriction enzyme digestion/ligation by the YE1 variant from plasmid YE1-BE3.
6. **YE1-BE3_SpCas9n**_**EQR**_ is a variant of the YE1-BE3_SpCas9n also made in our laboratory. The rAPOBEC1 deaminase sequence in plasmid pBK-EQR-BE3 was replaced by the YE1 sequence from plasmid YE1-BE3.
7. **YE1-BE3_SaCas9n**_**KKH**_ is a variant of the BE3_SaCas9n_KKH_ made in our laboratory. The rAPOBEC1 deaminase sequence in plasmid BE3_SaCas9n_KKH_ was replaced by the YE1 variant from plasmid YE1-BE3.
8. **BE4-Gam_SpCas9n**_**VQR**_ is a variant of the BE4-Gam (Addgene Inc. # 100806) made in our laboratory by PCR mutagenesis to create the VQR mutation in the SpCas9n gene.
9. **BE4-Gam_SpCas9n**_**EQR**_ is a variant of the BE4-Gam available at Addgene Inc. # 100806. This variant was made in our laboratory by PCR mutagenesis to create the EQR mutation in the SpCas9n gene.
10. **Target-AID SpCas9n**_**VQR**_ is a variant of the Target-AID enzyme described by Komor et al. (14). The original Target-AID enzyme contains the SpCas9n(D10A), a 105 amino acids linker, CDA1 (i.e., the activation induced cytidine deaminase (AID) that was modified by Nishida et al. (21)), a 11 amino acids linker and the UGI. The original plasmid, named nCas9-PmCDA1-UGI available at Addgene Inc. #76620 was modified by PCR mutagenesis to create the VQR version of the SpCas9n.
11. **Target-AID SpCas9n**_**EQR**_ is a variant of the Target-AID enzyme described by Komor et al. (14). The original Target-AID enzyme contains the SpCas9n(D10A), a 105 amino acids linker, CDA1 (i.e., the activation induced cytidine deaminase (AID) that was modified by Nishida et al. (21)), a 11 amino acids linker and the UGI. The original plasmid, named nCas9-PmCDA1-UGI and gRNA (HPRT)(Target-AID) available at Addgene Inc. #76620 was modified by PCR mutagenesis to create the EQR version of the SpCas9n.
12. **Target-AID SaCas9n**_**KKH**_ is a variant of the original plasmid called Target-AID available at Addgene Inc. #76620. The SpCas9n was replaced by SaCas9n_KKH_ with Gibson assembly. Briefly, the SpCas9n was removed by cutting the plasmid with two restriction enzymes Nhe1 and BsiW1.

### Co-transfection in HEK293T and SH-SY5Y cells of Cas9n/Cytidine deaminase plasmid and pBSU6 sgRNA

The transfection reagent (Lipofectamine 2000™) and Opti-MEM-1 ™ culture media) were purchased from Life Technologies Inc. (Carlsbad, CA). The HEK293T and SH-SY5Y cells were transfected with a plasmid coding for one Cas9n-deaminase and a sgRNA. The day before the transfection, 100,000 cells were seeded per well in a 24 well plate in DMEM (DMEM/F12 for SH) medium supplemented with 10% FBS and antibiotics (penicillin/streptomycin 100 μg/mL). The following morning, the culture medium was changed for 500 μl of fresh medium. The plate was maintained at 37°C for the time required to prepare the transfection solution. For the transfection, solutions A and B were first prepared. Solution A contained 48 μl of Opti-MEM™ and 2 μl of Lipofectamine™ 2000 for a final volume of 50 μl. Solution B was prepared as follows: a volume of DNA solution containing 800 ng of DNA was mixed (400ng of base editor plasmid and 400ng of pBSU6 sgRNA or GFP plasmid for negative controls) with a volume of Opti-MEM™ to obtain a final volume of 50 μl. Solutions A and B were then mixed by up and down movements and incubated at room temperature for 20 minutes. 100 μl of the mixed solution were then added to each well. The plate was let in the CO_2_ incubator for a period of 4 to 6 hours before a fresh medium change. The plate was incubated for 72 hours in the CO_2_ incubator before extraction of genomic DNA.

### Cell harvesting

Cells were detached 72 hours post-transfection by performing up and down movements in 1 ml culture medium with a pipette. These cells were transferred in an Eppendorf tube and centrifuged at 8000 RPM for 10 minutes. The supernatant medium was carefully removed without disturbing the cell pellets. These cell pellets were washed once with 1 ml of HBSS solution and centrifuged at 8000 RPM for 10 minutes. The HBSS was then carefully removed without disturbing the cell pellets.

### DNA extraction

The cells were lysed with 100 μl of lysis buffer containing 1% Sarkosyl and 0.5 M EDTA pH8 supplement with 10 μl of proteinase K solution (20 mg/ml). These tubes were incubated at 50^°^C for 15 minutes. 400 μl of 50 mM Tris pH8 were then added to each tube. Next, 500 μl of a mixture of phenol: chloroform: isoamyl alcohol (respectively 25:24:1) was added. The tubes were centrifuged at 16 000 RPM for 2 minutes. The aqueous upper phase was transferred to a new tube. 50 μl of NaCl 5 M were added to each tube and mixed thoroughly. One (1) ml of ice-cold ethanol 100% was added to each tube and mixed for genomic DNA precipitation. The tubes were centrifuged at 16000 RPM for 7 minutes and ethanol was carefully removed to avoid disturbing the DNA pellets. These DNA pellets were washed once with 400 μl ethanol 70%. The tubes were centrifuged at 16000 RPM for 5 minutes and ethanol was removed to permit to dry the DNA pellet rapidly by Speedvac vacuuming. The DNA was solubilized in 50-100 μl of sterile water and stored at −20°C until quantification was performed. The DNA solutions were dosed at 260 nm with a spectrophotometer.

### Stable SH-SY5Y APP cell lines generation

A lentivirus CMV-APP-P2A-Puromycin-WPRE was produced in a 10 cm petri dish containing 4 million HEK293T cells. 40 μg of four lentiviral plasmids were transfected (15 μg APP plasmid, 15 μg Gag-pol, 5 μg REV, 5 μg VSVG) with calcium phosphate method. The medium was replaced by 6 mL of fresh medium 16 hours later. The medium was harvested 72 hours post-transfection, filtered and directly poured on a 1 million SH-SY5Y plated 6-well. The medium was renewed the morning after. The cells were selected with 8 μg/mL of puromycin 72 h post-transduction.

### Supernatant analysis

At 72 hours post-transfection, 100 μL of the culture medium of the SH-SY5Y APP cell lines were harvested and filtered at 0.4 μm to remove cell debris. Protease inhibitors (1 mM PMSF + 1X complete tabs from Roche) were added. The media were then stored at −80°C. The concentrations of amyloid-β peptides 42 and 40 (most common AD biomarkers) were measured with Meso Scale Discovery Inc. (MSD, Rockville, MA) Neurodegenerative Disease Assay 6E10 kit. Standards and samples were prepared according to the manufacturer’s protocols and always tested in technical duplicates.

### Deep sequencing analysis

Deep sequencing samples were prepared by a PCR reaction with special primers containing a bar code sequence (BCS) sequences to permit the subsequent sequencing (i.e., BCS1: ACACTGACGACATGGTTCTACAGGTAGGCTTTGTCTTACAGTGTTA and BCS2: TACGGTAGCAGAGACTTGGTCTTGGTAATCCTATAGGCAAGCATTG). DNA sequences were analyzed with the Illumina sequencer. Roughly 10000 reads were obtained per sample.

### Bioinformatics analysis of the deep sequencing results

The proportion of wild-type versus edited genomes was estimated by counting the abundance of sequenced reads that contained exon 16 with the GCA wild-type codon (i.e., alanine) and the ACA edited codon (i.e., threonine).

## SUPPLEMENTAL INFORMATION

Supplemental Information includes 3 tables and 5 figures.

## AUTHOR CONTRIBUTIONS

A.G. designed the experiments, performed the experiments, and wrote the manuscript. J.R. assisted with the design of the experiments and provided technical assistance for molecular biology. F.G.B and T.B assisted for base-editors construction. G.L. revised and corrected the manuscript. J.P.T. conceived the experiments and corrected the manuscript.

## ACKNOWLEDGMENTS

This research project was supported by grants from the Weston Brain Institute. We also would like to thank the Quebec cell, tissue and gene therapy network &#8211;THÉCELL for his support. I would like to personally thank Dr. Serhat Gumrukcu for his advice and his revisions of this paper.

