## Supplemental tables and figures for "Base editing strategy allows insertion of the A673T mutation in APP gene to prevent the development of Alzheimer’s disease"

SUPPLEMENTAL RESULTS:

Table S1: List of sgRNA used in the study.

| | |
|---|---|
| SpCas9 VQR/EQR sgRNAs targeting sequences of 17 to 22 nucleotides tested: | |
| sgRNA 22bp | ATTCTGCATCCATCTTCACTTC |
| sgRNA 21bp | TTCTGCATCCATCTTCACTTC |
| sgRNA 20bp | TCTGCATCCATCTTCACTTC |
| sgRNA 19bp | CTGCATCCATCTTCACTTC |
| sgRNA 18bp | TGCATCCATCTTCACTTC |
| sgRNA 17bp | GCATCCATCTTCACTTC |
| SaCas9 sgRNAs tested: | |
| sgRNA 22bp | ATTCTGCATCCATCTTCACTTC |
| sgRNA 21bp | TTCTGCATCCATCTTCACTTC |
| sgRNA 20bp | TCTGCATCCATCTTCACTTC |
| sgRNA 19bp | CTGCATCCATCTTCACTTC |
| sgRNA 18bp | TGCATCCATCTTCACTTC |
| sgRNA 17bp | GCATCCATCTTCACTTC |

Table S2: Example of Deep-Sequencing analysis.

| | Target-AID-SpCas9nVQR 19 | BE3_SpCas9nVQR 19 |
|---|---|---|
| Total reads | 100% | 100% |
| Wild-Type | 31,8 | 66,0 |
| C1 | 3,8 | 1,0 |
| C2 | 3,0 | 0,3 |
| C3 | 0,1 | 3,6 |
| C4 | 0,3 | 0,0 |
| C5 | 0,8 | 0,3 |
| C1+C2 | 26,2 | 0,0 |
| C1+C3 | 0,2 | 2,1 |
| C1+C4 | 0,2 | 0,0 |
| C1+C5 | 0,2 | 0,0 |
| C2+C3 | 0,1 | 0,2 |
| C2+C4 | 0,1 | 0,0 |
| C2+C5 | 0,2 | 0,0 |
| C3+C4 | 0,1 | 0,2 |
| C3+C5 | 0,0 | 0,2 |
| C4+C5 | 0,0 | 0,0 |
| C1+C2+C3 | 2,0 | 1,0 |
| C1+C2+C4 | 1,6 | 0,0 |
| C1+C2+C5 | 4,2 | 0,0 |
| C2+C4+C5 | 0,0 | 0,0 |
| C2+C3+C5 | 0,0 | 0,0 |
| C1+C2+C3+C4 | 0,1 | 0,1 |
| C2+C3+C4+C5 | 0,3 | 0,2 |
| C1+C3+C4+C5 | 0,0 | 0,0 |
| C1+C2+C4+C5 | 0,0 | 0,1 |
| C1+C2+C3+C5 | 0,2 | 0,0 |
| C1+C2+C3+C4+C5 | 0,3 | 0,2 |
| Total | 75,60% | 75,70% |
| Mis-sequencing | 24,40% | 24,30% |

Table S3: Percentage of reduction of amyloid-β peptides 40 and 42 induced by the addition of the A673T mutation to wild type APP gene or to an APP gene containing the London mutation or containing a C1 deamination (E674K). The addition of the A673T mutation reduced the production of Aβ40 and Aβ42 peptides in all 3 situations.

| FAD mutation | Wild-Type | V717I (London) | A673T+E674K (C1+C2) |
|---|---|---|---|
| Abeta42 Decrease (%) | -46 | -65 | -53 |
| Abeta40 Decrease (%) | -63 | -81 | -44 |

Figure S1.

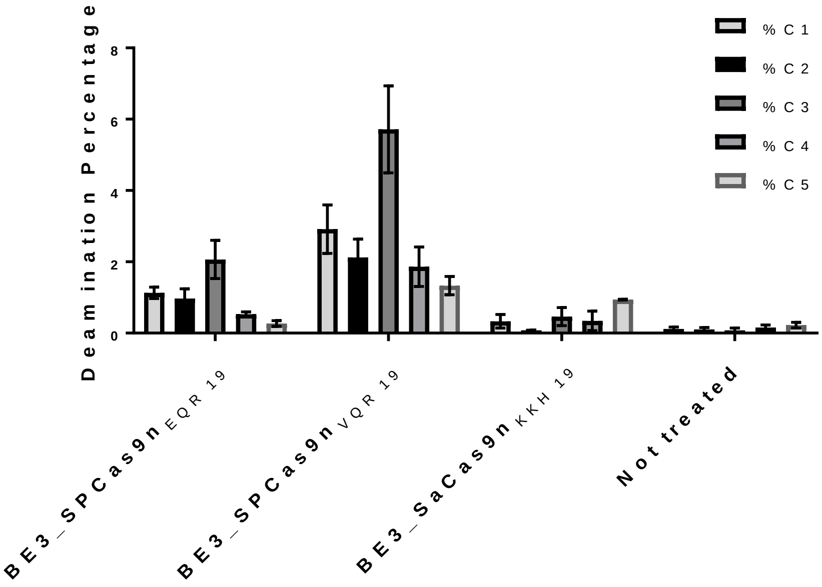


Figure S1: **Percentages of cytidine deamination produced by various enzymes and sgRNAs.** BE3_SpCas9nEQR, BE3_SpCas9nVQR, BE3_SaCas9nKKH enzymes test in SH-SY5Y cells. The figure illustrates the means +/- SEM (n=4).

Figure S2.

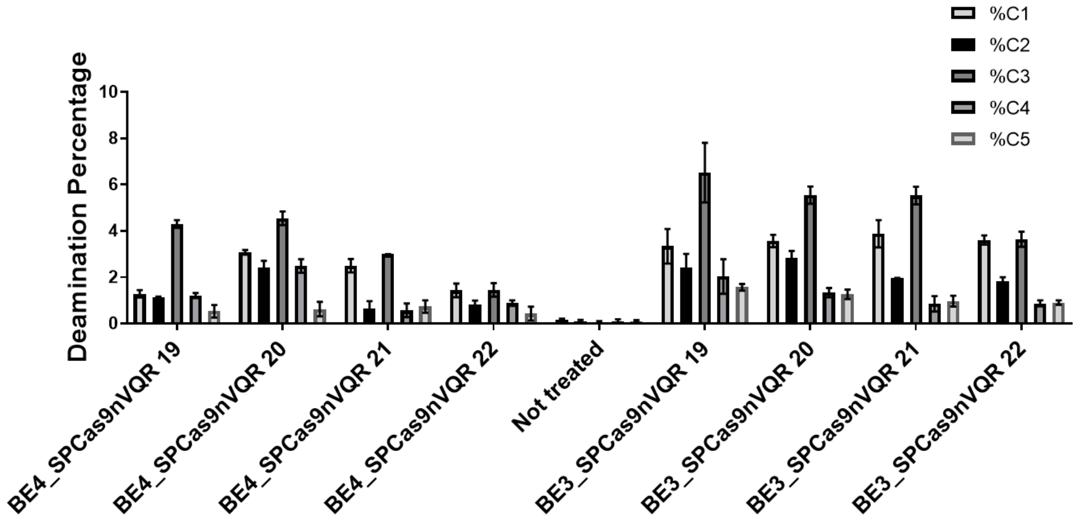

Figure S2: **Percentages of cytidine deamination produced by various enzymes and sgRNAs.** BE4_SpCas9nVQR and BE3_SpCas9nVQR enzymes test in SH-SY5Y cells. The figure illustrates the means +/- SEM (n=3).

Figure S3.

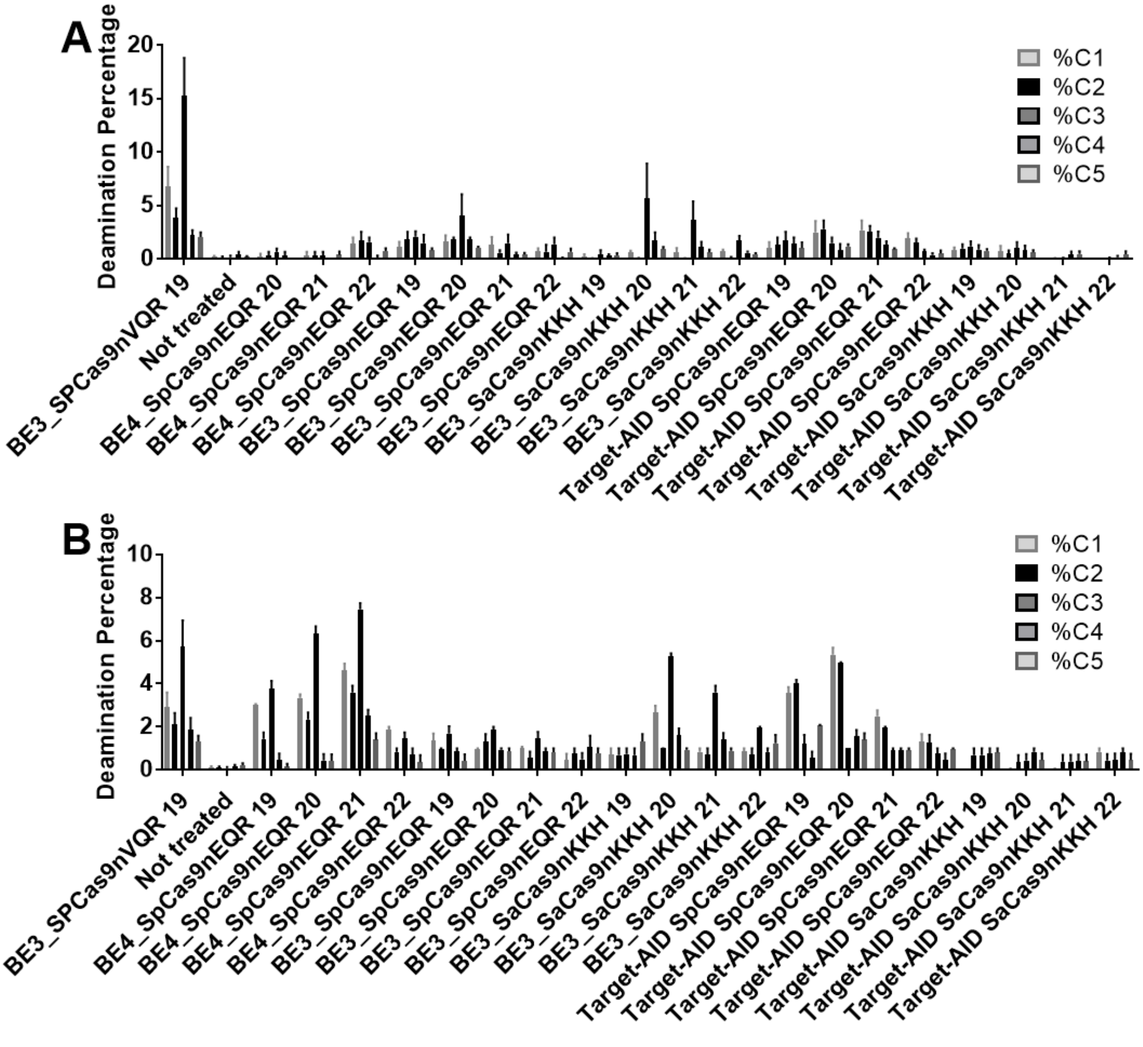


Figure S3: **Percentages of cytosine deamination produced by BE3_SpCas9nVQR, BE4_SpCas9nEQR, BE3_SpCas9nEQR, BE3_SaCas9nKKH, Target-AID_SpCas9nEQR, Target-AID_SaCas9nKKH enzymes.** In **A**, test in HEK293T cells. In **B**, test in SH-SY5Y. The figure illustrates the means +/- SEM (n=3).

Figure S4.

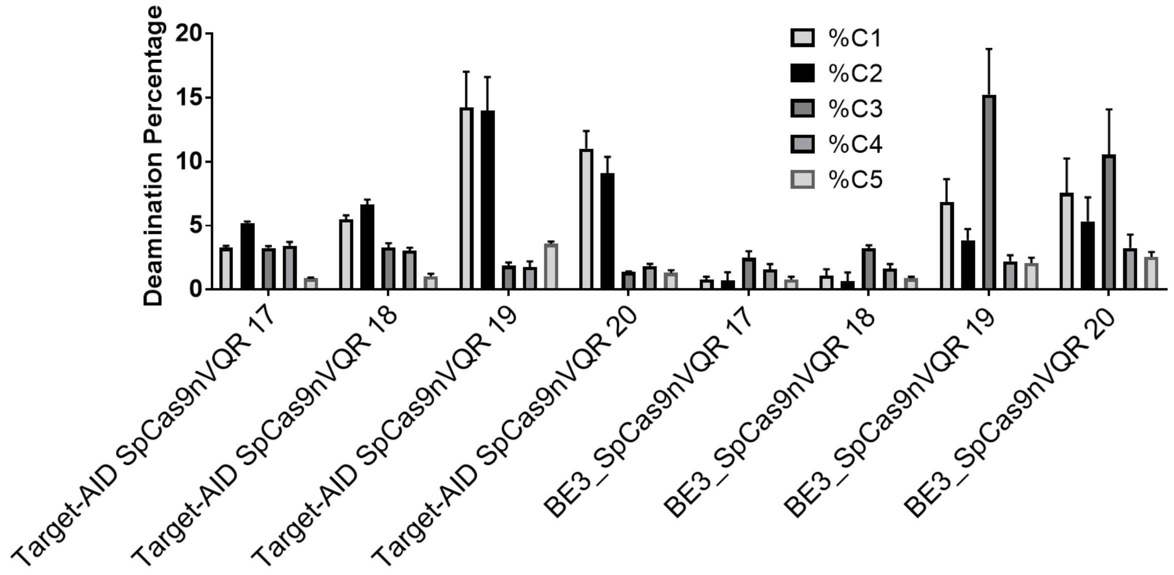


Figure S4: **Deamination efficiencies using various Cas9n-deaminases and sgRNAs targeting various numbers of nucleotides**. The difference of deamination in HEK293T cells of cytidines C1 to C5 produced by the Target-AID-SpCas9nVQR and BE3_SpCas9nVQR enzymes and two copies of a sgRNA targeting 17 to 20 nucleotides. The figure illustrates the means +/- SEM (n=3).

Figure S5

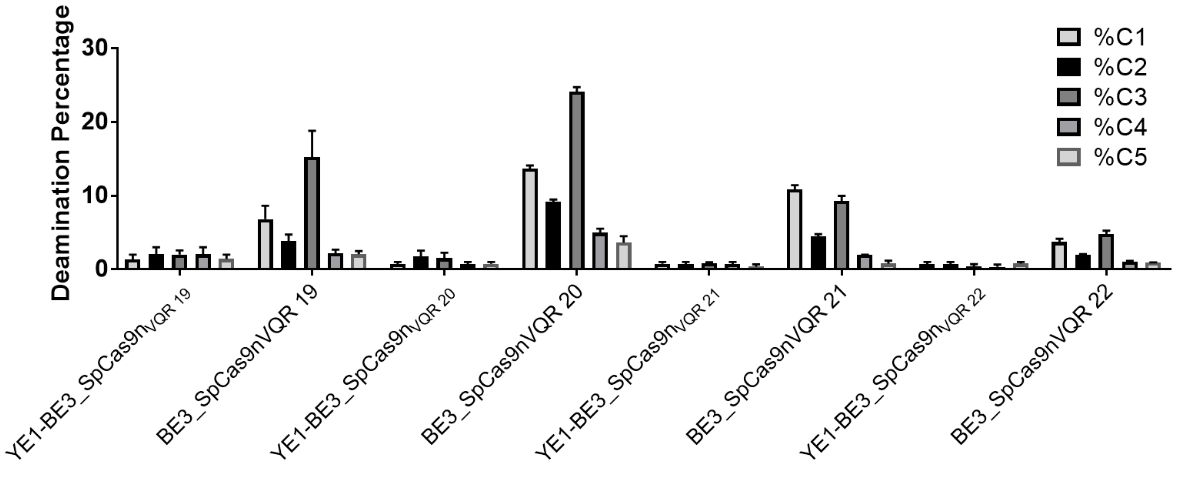


Figure S5: **Deamination efficiencies using various Cas9n-deaminases and sgRNAs targeting various numbers of nucleotides**. Difference between YE1-BE3_SpCas9nVQR and BE3_SpCas9nVQR in HEK293T cells. The figure illustrates the means +/- SEM (n=4).

Figure S6:

Off-Target Sites

Copy TSV

| Sequence | PAM | Score | Gene | Locus |
|---|---|---|---|---|
| CTGCATCCATCTTCACTTC | AGAG | 100.0 | APP (ENSG00000142192) | chr21:+25897633 |
| CTGAAGCCATCTTCACTTC | GGAG | 1.4 | | chr5:-74133250 |
| CTGCCTCCATCTTCACATG | TGAG | 0.5 | | chr11:+70758797 |
| CTGCTTCCAACTTCACTTT | GGAG | 0.5 | SPOCK2 (ENSG00000107742) | chr10:-72059638 |
| CAGGATCCATCTTAACTTC | TGAG | 0.4 | | chr12:-47654519 |
| CTGCTTCCATCTTCTGTTC | AGAG | 0.4 | | chr3:-36729648 |
| CTGCATCCTTCTCCACTTG | GGAG | 0.4 | | chr8:-67603274 |
| CTGAATCAATCTCCACTTC | AGAG | 0.4 | | chr12:-56572869 |
| ATCCATGCATCTTCACTTC | AGAG | 0.3 | | chr11:-20359838 |
| CTGCCCCCACCTTCACTTC | TGAG | 0.3 | | chr9:+85398673 |
| CTGCATCCATCTCTCCTTC | AGAG | 0.3 | | chr3:+176747607 |
| CTATTTCCATCTTCACTTC | AGAG | 0.3 | | chr9:-21442550 |
| ATGTATCCATCTTCACTGT | TGAG | 0.1 | | chr14:-35505959 |
| TTTCATCCATCTCCACTTT | AGAG | 0.1 | | chr5:-44542656 |
| TTTCATCCATCTTAACTAC | AGAG | 0.1 | | chr10:-120808163 |
| ATCCATCCACCTTCACTTG | TGAG | 0.1 | | chr4:+10179124 |

Figure S6: **Off target analysis performed with Benchling.com interface.**
